# Transdiagnostic brain module dysfunctions across sub-types of frontotemporal dementia: a connectome-based investigation

**DOI:** 10.1101/2023.10.29.564589

**Authors:** Zeng Xinglin, He Jiangshan, Zhang Kaixi, Xia Xiaoluan, Xu Shiyang, Zhen Yuan

## Abstract

**Background:** Frontotemporal dementia (FTD) is a complex neurodegenerative disorder encompassing heterogeneous subtypes, including behavioral variant frontotemporal dementia (BV-FTD), semantic variant frontotemporal dementia (SV-FTD), and progressive non-fluent aphasia frontotemporal dementia (PNFA-FTD). Unraveling the shared and distinctive brain module organizations among these subtypes is critical for unraveling the underlying neural basis of the disease. This study aims to explore brain module organization in FTD subtypes, seeking potential biomarkers and insights into their pathophysiology.

**Methods:** Resting-state functional magnetic resonance imaging data were obtained from the Frontotemporal Lobar Degeneration Neuroimaging Initiative, comprising 41 BV-FTD, 32 SV-FTD, 28 PNFA-FTD, and 94 healthy controls, following exclusion of participants with excessive head motion. Individual functional brain networks were constructed at the voxel level of gray matter and binarized with a 1% density threshold. Using predefined brain modules, we computed the modular segregation index (MSI) for each module, analyzed intermodular and intramodular connections to identify driving modular connections, and calculated the participation coefficient (PC) to detect regions with altered nodal properties associated with module integrity. A machine learning algorithm was employed for FTD subtype classification based on these matrices. Correlations between modular measures and clinical scores in each FTD subtype were examined.

**Results:** Distinct brain module organizations were observed across FTD subtypes, with lower MSI in the subcortical module (SUB), default mode network (DMN), and ventral attention network (VAN) in both BV-FTD and SV-FTD. Specifically, only BV-FTD exhibited disruption in the frontoparietal network (FPN). Notably, the bilateral fusional gyrus, left orbitofrontal cortex, left precuneus, and right insular thalamus showed significant group effects on PC, indicating altered nodal properties associated with module integrity. Our machine learning achieved a multiple classification accuracy of 85%. Correlations between specific network alterations and clinical variables in each FTD subtype were also identified.

**Conclusions:** These findings illuminate the diverse brain module organization in different FTD subtypes, offering insights into potential neurobiological differences that underlie the clinical heterogeneity of the disease. Regions with altered modular properties may serve as valuable biomarkers for early diagnosis and disease monitoring. Furthermore, understanding disruptions in modular connectivity provides valuable insights into the neuropathological mechanisms of FTD subtypes, paving the way for targeted therapeutic interventions.

## Introduction

Frontotemporal dementia (FTD) is a prevalent neurodegenerative syndrome primarily affecting the frontal cortex and temporal lobes [1]. FTD is the leading type of earliest onset dementia with an average age of onset around 60 years old [1, 2], also ranking as the third most common dementia type across all age groups [3]. FTD imposes significant socioeconomic burdens due to its high prevalence (3% to 26%) and relatively short disease duration of 7-8 years [3, 4]. FTD includes distinct subtypes: the behavioral variant of FTD (BV-FTD) marked by behavioral and executive function deficits, and primary progressive aphasia (PPA), which comprises the semantic variant of FTD (SV-FTD) and the progressive non-fluent variant of FTD (PNFA-FTD) [5-7]. BV-FTD is associated with impairments in behavioral and executive functions [5, 8]. Apart from sharing common features with BV-FTD in social and emotional symptom changes, like apathy, repetitive behaviors [3, 9], PPA refers to the gradual loss of linguistic skills within the initial two years of symptom onset [8]. To elaborate, SV-FTD is marked by semantic aphasia and associative agnosia, while patients with PNFA-FTD exhibit slow, labored, and halting speech production, retaining fluency but possibly displaying agrammatism [6].

As FTD remains a complex challenge for clinical treatment, understanding the neural mechanisms unique to each subtype and shared across them becomes paramount. Recent advances in magnetic resonance imaging (MRI) have provided insights into the neural underpinnings of various diseases, including FTD, by examining how clinical symptoms and behavioral attributes relate to functional network communication [10, 11]. The brain functions through synchronized interactions of multiple regions, giving rise to modular architecture—a network organization featuring specialized functional modules with dense intramodular connections and sparse intermodular connections [12, 13]. This modular organization is vital for balancing specialization and integration, supporting cognitive and behavioral functions [14]. Numerous studies reveal that disruptions in this modular connectivity, both within and between functional modules, are associated with neurodegenerative diseases like Alzheimer’s Disease (AD) [15], Parkinson’s disease [16], dementia with Lewy bodies [17], etc. Furthermore, the baseline brain modular architecture has been linked to cognitive decline in AD [18], and recovery in cognitive control function after intervention [13]. Given the strong correlation between symptoms and brain modular architecture in neurodegenerative diseases, it is valuable to find the common transdiagnostic and diagnosis-specific disorganization patterns across the disorders, like psychiatric disorders [19], neurodegenerative disease [16, 20, 21]. For instance, both BV-FTD and AD have exhibited global changes in modularity. BV-FTD, in particular, displayed a distinct disruption in the ventral attention network (VAN) compared to healthy controls and AD [20]. In contrast, PNFA-FTD has shown an increase in integration within the speech production network [22]. Notably, there is a lack of previous research exploring the modular architecture of functional networks in SV-FTD. While Ramanan, et al. [23] conducted a transdiagnostic comparison between BV-FTD and SV-FTD, they found that both subtypes exhibited comparable magnitude of atrophy to frontal regions, with only SV-FTD displaying severe temporal lobe atrophy. It’s important to note that previous studies have primarily focused on functional connectivity within brain regions and other graph theory properties in resting-state MRI [9, 24].

Consequently, direct evidence regarding common transdiagnostic and diagnosis-specific patterns of disorganization in brain modularity across FTD subtypes remains limited.

This study seeks to delve into the modular architecture of high-resolution functional brain networks at the voxel-wise level, utilizing resting-state fMRI data from individuals with Frontotemporal Dementia (FTD). We anticipate that patients with distinct dementia subtypes will manifest both shared and subtype-specific alterations in terms of module-specific segregation and integration, marked by an increased number of intermodular connections. We also aim to identify the brain regions that govern these connectivity patterns. To facilitate subtype classification within the FTD spectrum, we harness machine learning techniques. Machine learning, tailored for intricate multivariate data, holds promise as an adjunct in the diagnostic process by offering decision support[25]. Previous studies have use machine learning algorithms to classify between the subtypes of dementia [25-27] and subtypes of FTD [28, 29] in structural MRI. The accuracy of machine learning model between subtypes of FTD was ranged from 0.7-0.94. Our study distinguishes itself by utilizing features derived from resting-state fMRI, presenting an alternative method. Finally, our investigation extends to examining the association between modular architecture, clinical symptoms, and the rate of cognitive decline in FTD.

## Method

### Participants and Image Acquisition

This study drew its participant pool from the Frontotemporal Lobar Degeneration Neuroimaging Initiative (FTLDNI) databases. The final dataset consisted of 94 HCs, 42 individuals with BV-FTD, 31 with SV-FTD, and 28 with PNFA-FTD. Inclusion criteria required participants to have 3T high-quality resting state fMRI and T1-weighted MRI scans, as well as a stable diagnostic label of HC, BV-FTD, SV-FTD, or PNFA-FTD (with no discrepancies between imaging label and demographic data). A total of 20 participants were excluded due to excessive head motion exceeding 3 mm or 3°. Diagnostic criteria for BV-FTD were adhered to according to references [5, 30], and the diagnosed criteria of SV-FTD and PNFA-FTD were followed references [5, 6].

Image acquisition took place at three centers using Siemens TrioTim 3-T scanners. T1-weighted MR images were obtained using the MP-RAGE sequence with the following parameters: repetition time (TR)=2300 ms, echo time (TE)=3 ms, flip angle=9.0°, matrix size=240×256, slice number=160, voxel resolution: 1 □× 1 □× □1 mm. The resting state fMRI scan lasted for 8 minutes, resulting in 240 volumes. The image acquisition parameters were as follows: TR=2000 ms, TE=27 ms, flip angle=80°, voxel resolution: 2.5 □× □2.5 □× □3 mm, and slice number=48.

### Data Preprocessing and Network Construction

The preprocessing of fMRI data was carried out using fMRIPrep with default parameters. The resultant BOLD time series underwent detrending and deconfounding, involving the removal of 18 variables, including the six estimated head-motion parameters (tran_x_, tran_y_, trans_z_, rot_x_, rot_y_, rot_z_), the first six noise components computed using anatomical CompCorr, and six DCT-basis regressors using nilearn’s clean_img pipeline with default parameters. Subsequently, all fMRI data were resampled to 3-mm isotropic voxels and spatially smoothed with a 6-mm Gaussian kernel.

Individual functional networks were constructed at the voxel level within a gray matter (GM) mask based on the Brain Connectome atlas, which comprised 41,423 voxels corresponding to cerebral regions. By calculating Pearson’s correlation coefficients between all pairs of GM voxels, a 41,423 × 41,423 correlation matrix was generated for each subject. These individual correlation matrices were then binarized using a density threshold of 1%, resulting in 17,158,253 retained edges with the strongest positive correlation strength.

**Figure 1.**
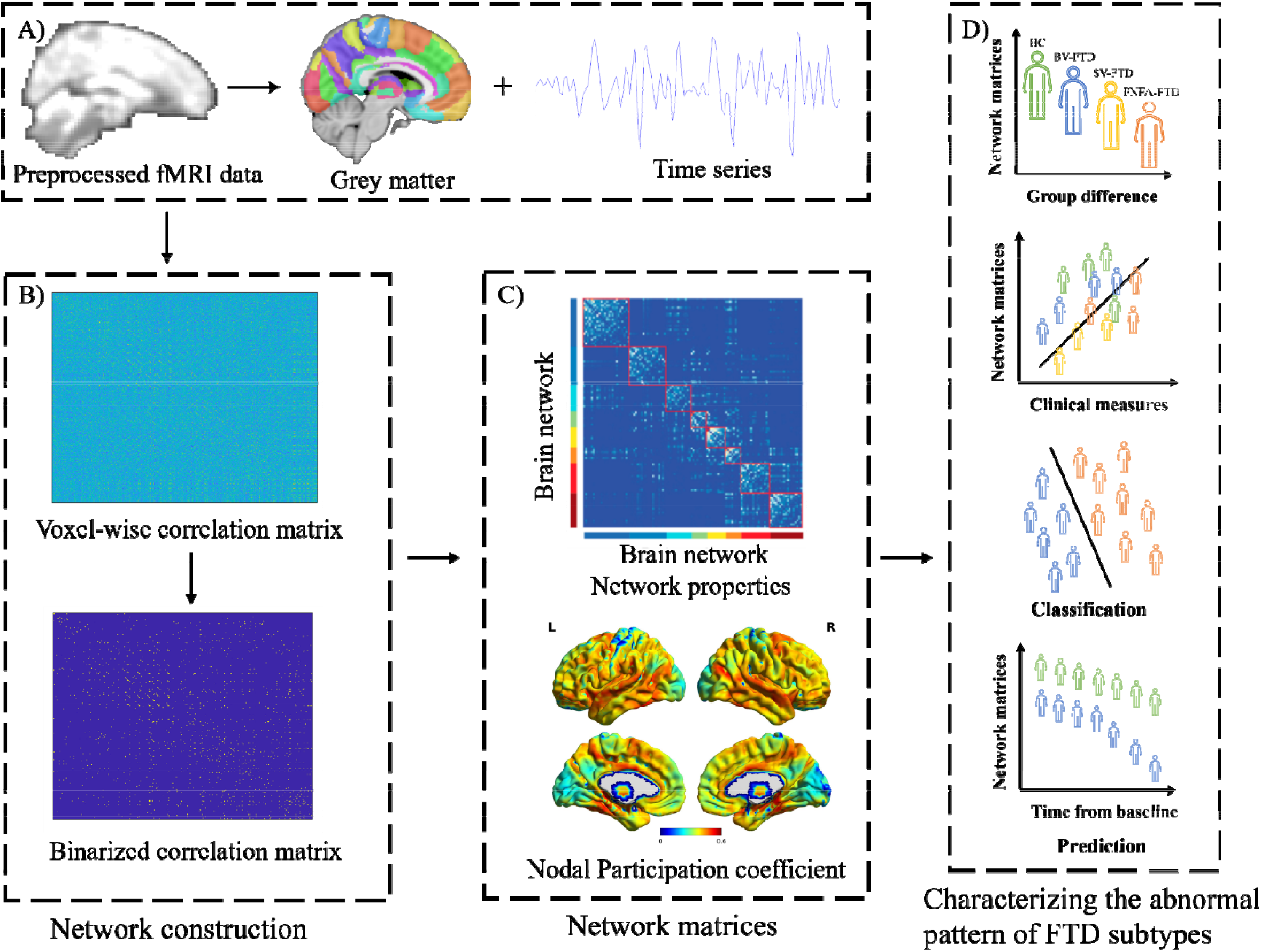
Data analysis pipeline overview. A) Data preprocessing was conducted, and gray matter time series were extracted using the Brainnetome Atlas. B) Voxel-wise correlation matrices were generated by computing Pearson correlation coefficients for each voxel, followed by binarization based on connectivity strength. C) Network matrices were calculated, incorporating module-specific segregation, intra- and intermodular connections, and nodal participation coefficients. D) Group differences in network metrics were compared. Additionally, the study explored the correlation between clinical measures, disease progression, and network metrics. Artificial intelligence was utilized to classify and predict FTD subtypes.

### Modular Architecture

In analyzing the modular architecture of individual brain networks, we followed this procedure: We partitioned whole-brain functional networks into eight modules using a predefined cortical parcellation to reduce disorder-related biases. These modules included the Frontoparietal Network (FPN), Default Mode Network (DMN), Dorsolateral Attention Network (DAN), VAN, Limbic Network (LIM), Visual Network (VIS), Sensorimotor Network (SSN), and a Subcortical Module (SUB) encompassing structures like the thalamus, putamen, hippocampus, caudate, amygdala, and pallidum. For each module (i), we calculated the Modular Segregation Index (MSI) as follows:

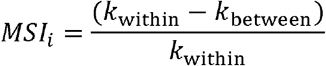

where k_within_is the number of intramodular connections of module i, and k_between_ is the total number of intermodular connections between module i and all other modules. Higher MSI values indicate greater functional segregation, while lower values suggest greater functional integration. We further determined the specific modular connections responsible for driving module specialization by computing intermodular connections for each pair of modules and intramodular connection numbers for individual modules. Finally, we calculated the Participation Coefficient (PC) of each voxel to identify regions with altered nodal properties associated with module integrity.

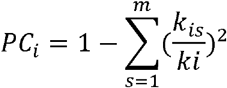

where m is the number of modules, k_is_ is the number of edges between node i and other nodes in module s, and k_i_ is the total number of edges linking to node i. Regions with high PC values are regarded as network connectors capable of communicating within their module and with other modules. PC was calculated based on individual modular partitions, and PC maps were compared among the four groups. For regions displaying significant group effects on PC, we conducted a region-to-module connectivity analysis to determine the contributions of different module connections to group differences in PC.

### Machine Learning

The modular architecture, including network and node matrices, was employed to train machine learning models for classifying FTD subtypes and HCs. Initially, all data were normalized to a range between 0 and 1 based on their minimum and maximum values. Given the relatively small sample size and the large number of features, kernel Principal Component Analysis (PCA) was used to reduce the dimension of the feature space. After feature selection, three established supervised machine learning methods—decision trees, Naïve Bayesian, and random forests—were employed to construct the classifier.

Given the limited sample size, 10-fold cross-validation was employed to enhance the model’s generalization ability. In this approach, 90% of the data served as the training dataset to train the model, while the remaining 10% constituted the test dataset, and this process was repeated 10 times. Four performance metrics, including accuracy, sensitivity, specificity, F1 score, and the area under the curve (AUC), were used to assess the classifier’s performance in distinguishing the four groups.

### Statistical Analysis

To assess group differences in demographic and clinical characteristics, we employed one-way ANOVA or the chi-squared test, followed by post hoc analyses where necessary. Nonparametric permutation tests were utilized for the modular metrics. Specifically, for each metric, we initially calculated the Freal-statistic, which quantifies the ratio of between-group variance to within-group variance, to estimate the group effect. We then created an empirical null distribution of the group effect by randomly redistributing all participants into four groups with the same sample sizes as the actual categories. Subsequently, we recomputed the group effect, F_surrogate_, among the reorganized groups through 10,000 permutations. This allowed us to calculate a P-value by determining the proportion of the observed group effect Freal within the null distribution. Post hoc analyses for metrics exhibiting a significant group effect were conducted using nonparametric permutation tests in a similar manner. Here, we estimated the t-statistic to assess between-group differences within the permutations (10,000 permutations). We also incorporated age and gender as control variables in the permutation analyses, and additional analyses were conducted considering smoking history as a covariate.

For module-specific metrics, including MSI and intra- and intermodular connections, we applied false discovery rate (FDR) correction to account for multiple comparisons. Specifically, for network metrices, this correction spanned right MSI, eight intramodular and 28 intermodular connections, and significance was defined as a corrected P-value of < .05. In the case of PC, we initially established a height threshold of P < .01 for each voxel and recorded the maximal cluster size exceeding this threshold in each permutation test. Following 10,000 permutations, we observed a null distribution of cluster sizes and determined the significance threshold for cluster correction as the 95th percentile of this distribution, corresponding to P < .05. Importantly, the significance level for post hoc pairwise comparisons was set to an FDR-corrected P-value of < .05 across all group pairs.

Finally, Spearman’s correlation analyses were conducted to explore relationships between each module-related measure and clinical variables. An FDR correction was applied across clinical variables, and significance was established at a corrected P-value of < .05. Age and gender were controlled for in the correlation analyses. Notably, these correlations were conducted across all patients for measures showing transdiagnostic alterations and within each patient group for diagnosis-specific alterations. It should be mentioned that because most participants in the HC groups lacked clinical dementia ratings, FAQ scores, the correlation of CDR was measured in FTD groups, while other clinical measures were analyzed across all groups. Additionally, to investigate the associations between module-related measures and longitudinal changes in cognition among FTD subtypes, Spearman correlations were performed. The degree of longitudinal cognitive changes was quantified using a linear regression model, and the coefficient estimates were correlated with module-related measures while controlling for age and gender as covariates. It is noted that all reported p values were corrected by FDR.

## Results

### Demographic and Clinical Characteristics

No significant gender or education differences were observed among the four groups. However, the age of participants in the PNFA-FTD group was significantly higher compared to the other three groups (Table 1). In terms of clinical measurements, the groups exhibited variations in the following assessments: Clinical Dementia Rating (CDR_total), CDR_box, CDR_language, CDR_behaviors, Mini Mental State Exam (MMSE), Montreal Cognitive Assessment (MoCA), California Verbal Learning Test (CVLT), Geriatric Depression Scale (GDS), Functional Activities Questionnaire (FAQ), Interpersonal Reactivity Index (IRI), Behavioral Activation Scale (BAS), and Behavioral Inhibition Scale (BIS).

**Table 1.** demographic and clinical data of FTD subtypes.

| Item | HC | BV-FTD | SV-FTD | PNFA-FTD | Statistics |
| --- | --- | --- | --- | --- | --- |
| Age | 62.87 (7.46) | 61.07 (6.277) | 62.84 (6.367) | 68.48 (14.171) | $F(3, 193) = 6.433, p = 0.0003$ |
| Gender (male/female) | 42/54 | 28/14 | 19/12 | 14/14 | $\chi^2 = 7.35, p = 0.06$ |
| Education | 21.70 (18.31) | 19.49 (18.47) | 19.19 (15.04) | 19.00 (15.61) | $F(3, 193) = 0.31, p = 0.81$ |
| CDR_total | 0.04 (0.14) | 1.11 (0.59) | 0.64 (0.32) | 0.43 (0.42) | $F(3, 155) = 63.84, p < 0.0001$ |
| CDR_box | 0.06 (0.18) | 6.23 (3.09) | 3.69 (2.08) | 1.95 (2.17) | $F(3, 155) = 79.99, p < 0.0001$ |
| CDR_language | 0 | 0.74 (0.55) | 1.02 (0.57) | 1.30 (0.68) | $F(3, 149) = 59.56, p < 0.0001$ |
| CDR_behaviors | 0.01 (0.07) | 1.61 (0.76) | 1.08 (0.63) | 0.40 (0.39) | $F(3, 149) = 84.87, p < 0.0001$ |
| MMSE | 29.3 (0.86) | 24.17 (4.57) | 25.17 (4.89) | 25.84 (3.99) | F (3, 188) = 29.65, p < 0.0001 |
| MoCA | 27.72 (1.49) | 19.86 (6.70) | 19.07 (4.27) | 22.7 (3.56) | F (3, 84) = 26.7, p < 0.0001 |
| CVLT | 29.9 (3.78) | 21.41 (6.99) | 17.57 (6.69) | 23.32 (5.93) | F (3, 132) = 28.11, p < 0.0001 |
| GDS | 3.74 (4.38) | 7.81 (6.90) | 8.65 (5.63) | 6.00 (4.34) | F (3, 170) = 9.139, p < 0.0001 |
| FAQ | 0.02 (0.14) | 16.95 (7.23) | 9.11 (5.76) | 6.48 (7.17) | F (3, 136) = 75.54, p < 0.0001 |
| BAS | 15.54 (8.38) | 16.11 (4.62) | 18.18 (3.72) | 17.77 (3.23) | F (3, 83) = 0.583, p = 0.628 |
| BIS | 29.25 (16.58) | 33.03 (7.37) | 31.62 (5.37) | 32.78 (6.21) | F (3, 83) = 1.232, p = 0.304 |

CDR: Clinical Dementia Rating; MMSE: Mini Mental State Exam; MoCA: Montreal Cognitive Assessment; CVLT: California Verbal Learning Test; GDS: Geriatric Depression Scale; FAQ: Functional Activities Questionnaire; IRI: Interpersonal Reactivity Index; BAS: Behavioral Activation Scale; BIS: Behavioral Inhibition Scale.

### Module Segregation of Brain Networks in subtypes of FTD Transdiagnostic Alterations

Significant group effects on the MSI (Table S1, Figure 2 Panel B) were observed in the SUB (F(3,193) = 9.5464, P < .0001), DMN (F(3,193) = 10.062, P < .0001), FPN (F(3,193) = 9.158, P < .0001), VAN (F(3,193) = 13.85, P < .0001). The post hoc analysis revealed a common decreased MSI of SUB, DMN, VAN in both BV-FTD and SV-FTD, compared to HC. Meanwhile, only BV-FTD was found decreased MSI in FPN compared to HC. However, compared to HC, no significant difference in MSI have been found in PNFA-FTD. Further analyses revealed that these alterations were mainly driven by increased numbers of intermodular connections (Table S1, Figure 2 Panel C-D), including SUB-SSN (F(3,193) = 5.928, P = .008), SUB-VAN (F(3,193) = 5.059, P = .013), SUB-DMN (F(3,193) = 4.666, P = .014), DMN-VAN (F(3,193) = 4.91, P = .013), FPN-AN (F(3,193) = 6.318, P = .007), FPN-DAN (F(3,193) = 4.665, P = .014), FPN-AN (F(3,193) = 5.72, P = .008), AN-VAN (F(3,193) = 5.72, P = .0008), AN-SSN (F(3,193) = 4.12, P = .0262), within VAN (F(3,193) = 3.97, P = .0288), within SSN (F(3,193) = 4.978, P = .0024), SSN-VN (F(3,193) = 9.761, P < .0001).

**Figure 2.**
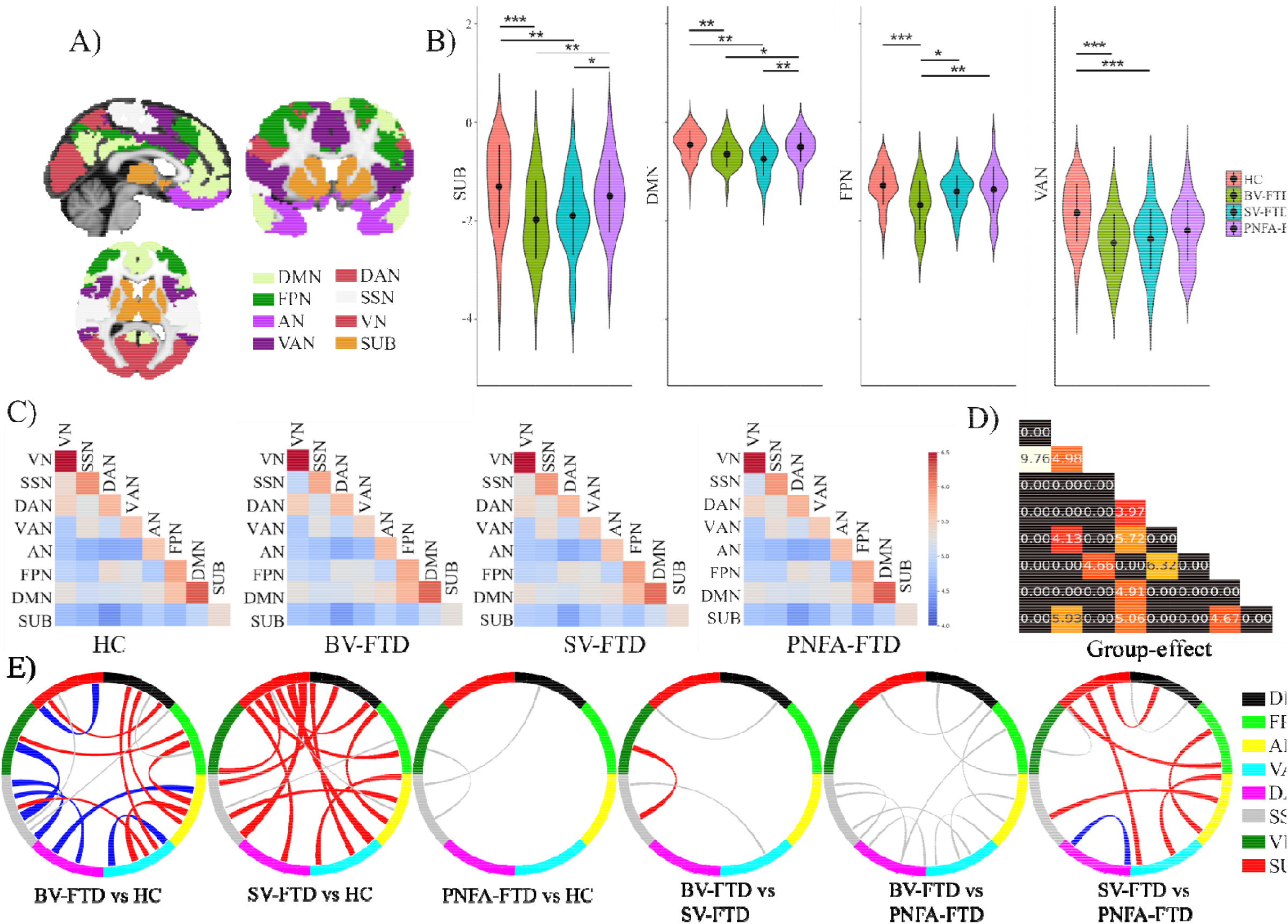
Differences in module segregation index and intra- and intermodular connections among the four groups A) An 8-module parcellation was generated from seven resting state networks and subcortical regions. B) Between-group differences in module segregation in VAN, FPN, DMN, SUB. C) Matrices display intra- and intermodular connectivity strength for each group. D) The heatmap shows group effects of intra- and intermodular connectivity among the groups. E) Between-group differences in intra- and intermodular connections for each pair of groups, with blue representing decreased connectivity, red representing increased connectivity after FDR correction, and gray indicating group differences that were not significant after FDR correction. ^* * *^, P <0.001, ^* *^, P<0.01, ^*^, P<0.05.

### Diagnosis-Specific Alterations

Compared to PNFA-FTD, BV-FTD exhibited dereased MSI in SUB (T(2, 67) = -3.829, P = 0.001), DMN (T(2, 67) = -2.777, P = 0.03), and FPN (T(2, 67) = -3.926, P = 0.0007). In comparison to SV-FTD, BV-FTD showed decreased MSI in FPN (T(2, 70) = -3.022, P = 0.015). SV-FTD displayed decreased MSI in SUB (T(2, 58) = -2.982, P = 0.017) and DMN (T(2, 58) = -3.741, P = 0.001) compared to PNFA-FTD. However, no intra- or inter-modules differences were found between BV-FTD and PNFA-FTD after controlling for the FDR (Figure 2 Panel E). Additionally, compared to PNFA-FTD, SV-FTD exhibited altered connectivity in SUB-DMN, SUB-FPN, SUB-VAN, FPN-AN, AN-VAN, AN-SSN, and within DAN (Figure 2 Panel E).

### Roles of Nodes in the Modular Brain Networks in subtypes of FTD

#### Transdiagnostic Alterations

Figure 3A illustrates the average Participation Coefficient (PC) maps in the brain networks for each group. A significant group effect on PC was observed in the bilateral fusiform gyrus (FG), left orbitofrontal cortex (OFC), left precuneus, and right insular (Table S1, Figure 3 B-C). When compared to HC, only BV-FTD was found lower PC values in right fusiform gyrus, left OFC, and left precuneus. Additionally, lower PC values in the right fusiform gyrus were observed in both BV-FTD and PNFA-FTD. Intriguingly, all subtypes of FTD displayed higher PC values in the right insular compared to HC. Specifically, these alterations in network nodes were mainly attributed to the greater number of connections (Figure 3 D) involving right insular-VAN (F(3,193) = 11.306, P < 0.0001), right insular-FPN (F(3,193) = 5.449, P = 0.008). Meanwhile, compared to HC, the altered PC values of left OFC contributing to the greater number of connectivity in left OFC-DMN (F(3,193) = 8.454, P = 0.0002), left OFC-FPN (F(3,193) = 5.784, P = 0.002), left OFC-AN (F(3,193) = 5.784, P = 0.002), left OFC-VAN (F(3,193) = 4.941, P = 0.005).

**Figure 3.**
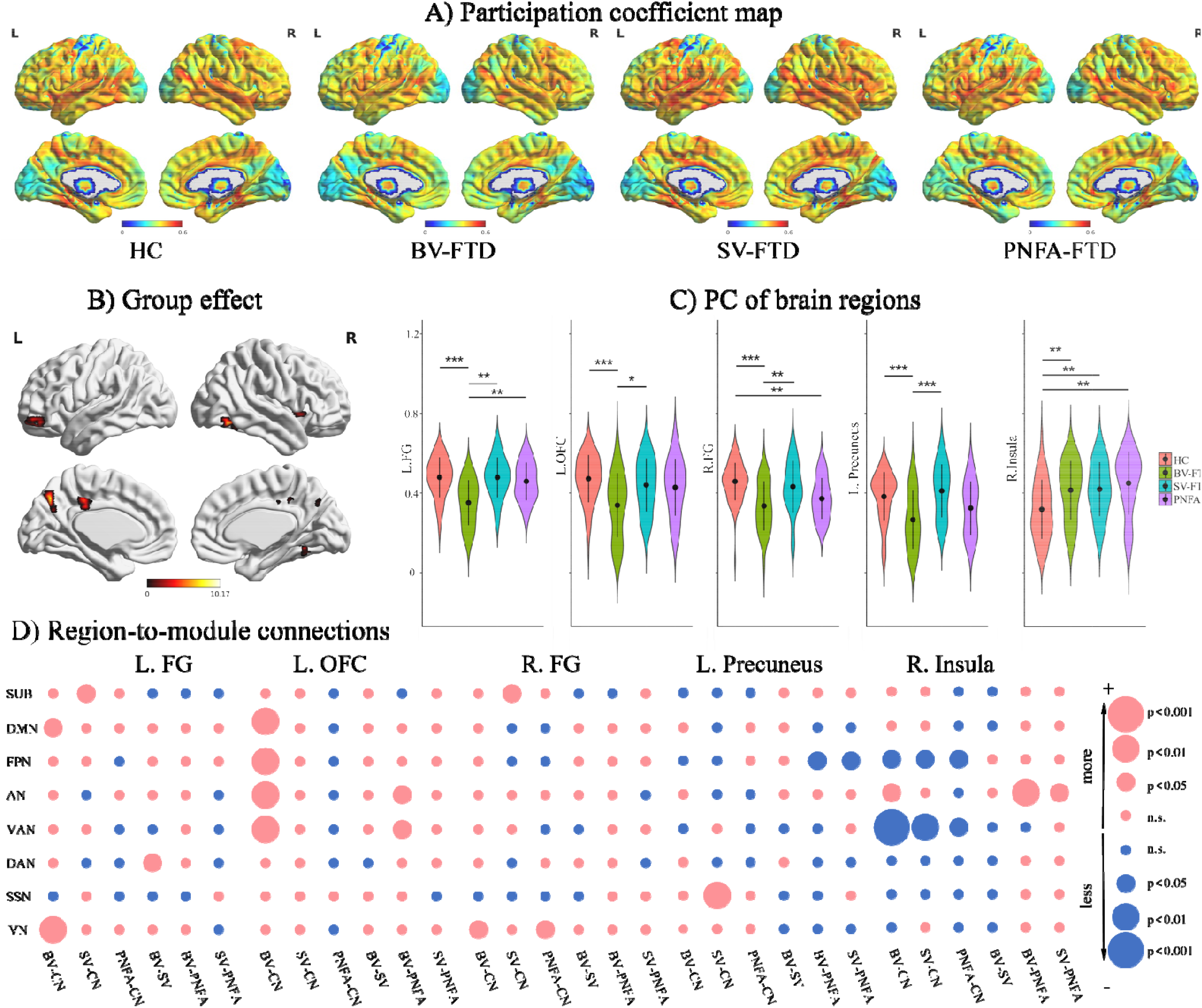
PC values and connections across FTD subtypes. A) Mean PC map for each group. B) Regions showing significant group effects on PC. C) Pairwise comparisons in regions with significant group effects on PCs. D) Between-group differences in region-to-module connections between each pair of groups. The blue representing decreased connectivity while red presenting increased connectivity after FDR correction. ^* *^, P <0.001, ^* *^, P<0.01, ^*^, P<0.05.

#### Diagnosis-Specific Alterations

In comparison, patients with BV-FTD exhibited significantly lower PC values in left FG (T(2, 58) = - 4.406, P = 0.001), left OFC (T(2, 58) = -3.077, P = 0.012), right FG (T(2, 58) = -3.73, P = 0.001) left precuneus (T(2, 58) = -4.735, P < 0.001) than patients with SV-FTD. The BV-FTD also had lower PC values in left fusiform gyrus than PNFA-FTD (T(2, 58) = -5.825, P < 0.001). No group difference of PC values has been found between SVFTD and PNFAFTD. Specifically, compared to SVFTD, the BV-FTD was found increased connectivity in left FG-VN, while compared to PNFA-FTD, the BV-FTD was found increased connectivity in left OFC-AN, left OFC-VAN, right insular-AN, and decreased connectivity in left precuneus-FPN. Compared to PNFA-FTD, the SV-FTD was found increased connectivity in right insular-AN and decreased connectivity in left precuneus-FPN.

### Relationship Between Brain Module Measures and Clinical Variables

The MSI of various brain networks exhibited significant correlations with clinical variables (Figure 4). Regarding general cognitive function, the MSI of the SUB (r = -0.32, P = 0.009), FPN (r = -0.34, P = 0.004), and VAN (r = -0.27, P = 0.02) were inversely correlated with CDR total scores, suggesting that higher CDR scores corresponded to more pronounced cognitive impairment. No significant correlations were found between MSI values and CDR language scores, but both the MSI of SUB (r = -0.34, P = 0.005) and FPN (r = -0.35, P = 0.005) showed negative associations with CDR behavior scores. In the context of episodic verbal learning and memory, as measured by the CLTV score, a positive relationship was observed with the MSI of SUB (r = 0.33, P = 0.002) and VAN (r = 0.24, P = 0.02). Additionally, higher levels of depression were negatively associated with the MSI of the DMN (r = -0.24, P = 0.009) and VAN (r = -0.26, P = 0.004). Functional activity, as assessed by the FAQ scores, displayed negative correlations with the MSI values of SUB (r = -0.31, P = 0.01), FPN (r = -0.4, P = 0.003), and VAN (r = - 0.36, P = 0.005). Furthermore, when examining the relationship between MSI values and the rate of cognitive decline, the MSI of SUB (r = -0.31, P = 0.013), FPN (r = -0.3, P = 0.018), VAN (r = -0.31, P = 0.013), DAN (r = -0.3, P = 0.018) were found to be negatively associated with the beta values of CDR total scores.

**Figure 4.**
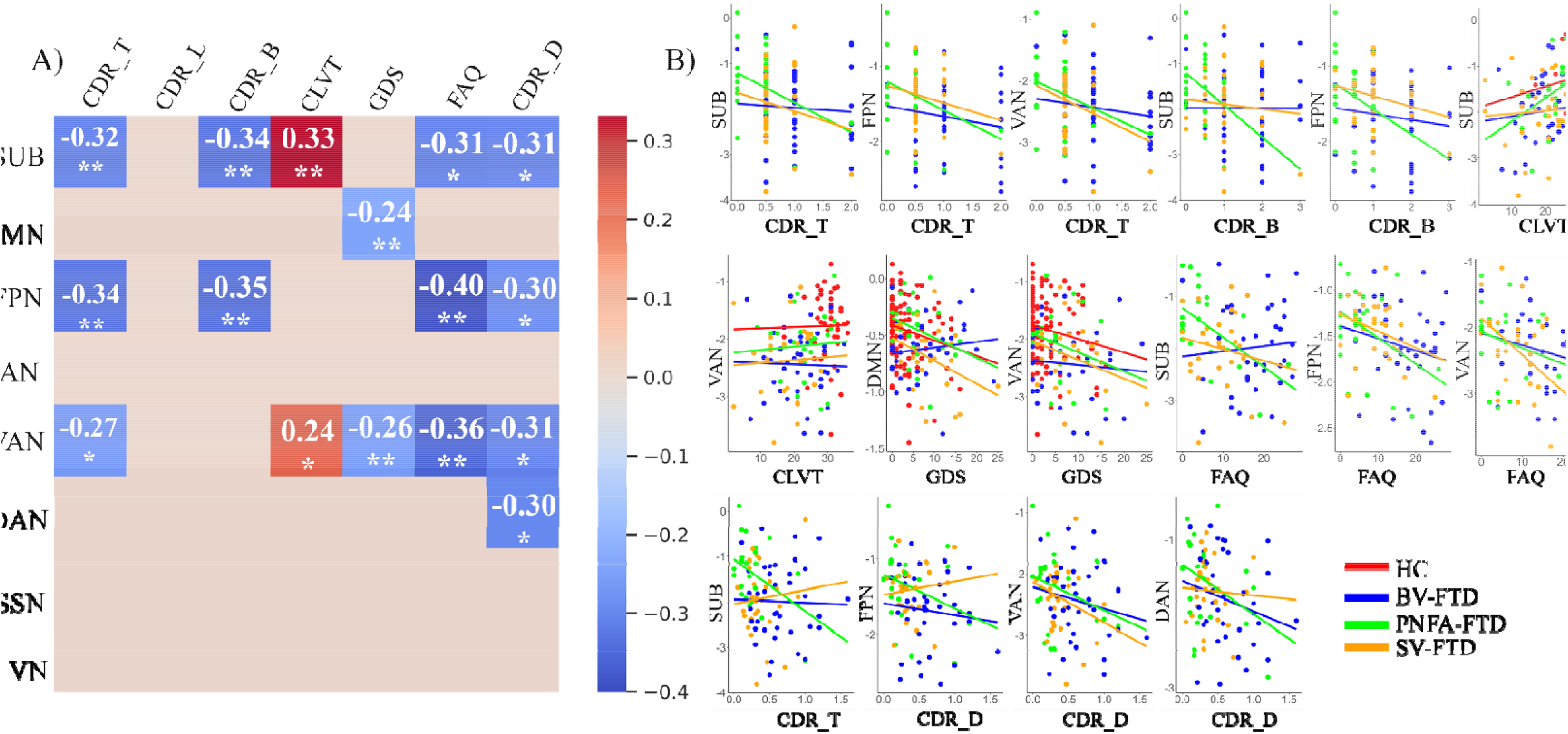
Correlations between clinical variables and MSI of networks. A) The heatmap between clinical variables and MSI of networks. B) The scatter plot between clinical variables and MSI of networks in each group. ^* * *^, P <0.001, ^* *^, P<0.01, ^*^, P<0.05.

### Machine learning performance

A machine learning approach using naïve Bayesian classification was employed to categorize individuals into the four groups based on features extracted from network matrices and PC values. The model achieved an overall accuracy of 85% (Figure 5). When considering HCs, the classification accuracy reached 90%, with a precise rate of 82%, a recall rate of 82%, and an AUC value of 0.849. For BV-FTD, the classification demonstrated a precise rate of 78%, a recall rate of 100%, and an AUC value of 0.941. PNFA-FTD classification yielded a precise rate of 62%, a recall rate of 62%, and an AUC value of 0.864. Finally, SV-FTD classification was highly precise at 92%, with a recall rate of 100% and an AUC value of 0.902.

**Figure 5.**
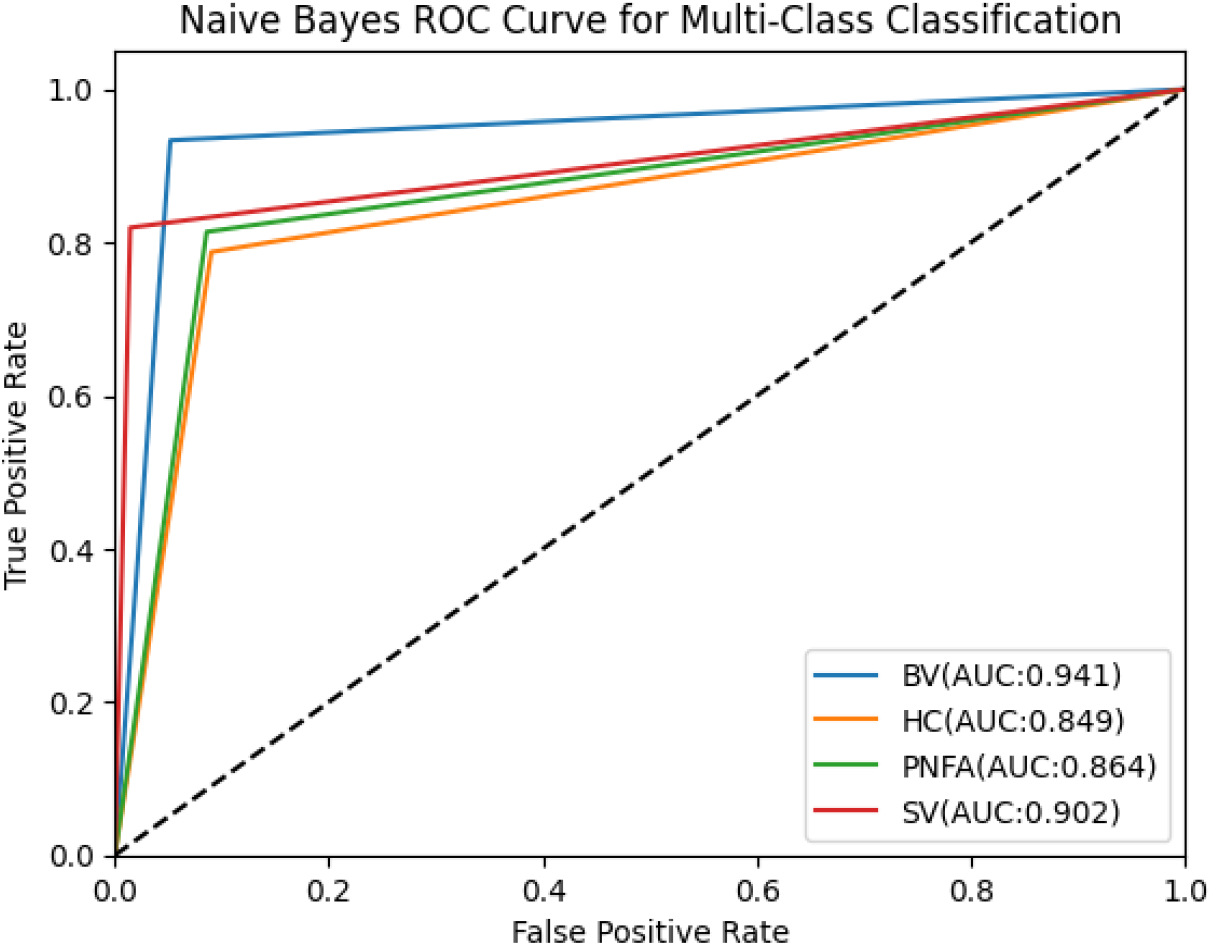
Performance of multiple classification.

## Discussion

Using a high-resolution connectomic analysis framework, this study identified both transdiagnostic and diagnosis-specific alterations in modular brain networks among different subtypes of FTD. Notably, primary transdiagnostic alterations included reduced modular specialization in the SUB, DMN, FPN, and VAN, driven by an increased number of intermodular connections. Additionally, specific nodes in the FG, OFC, precuneus, and insular cortex were found to be altered. On the other hand, diagnosis-specific findings were more pronounced in BV-FTD, with broader alterations in modular architectures compared to SV-FTD and PNFA-FTD. These findings provide crucial insights into the transdiagnostic and diagnosis-specific pathophysiological mechanisms of different FTD subtypes from a modular perspective.

### Greater Transdiagnostic Integration Among Modules

The modular architecture of the brain represents a highly efficient network organization that optimally balances specialization and integration, ensuring an equilibrium between energy consumption and communication efficiency. Notably, our investigation revealed heightened integration in SUB, DMN, FPN, and VAN in both BV-FTD and SV-FTD patients. This suggests a trend towards dedifferentiation in the overall brain network organization within these individuals. Previous research has reported lower average clustering coefficients, decreased global efficiency, and increased path lengths in both BV-FTD and SV-FTD cases [31]. Global efficiency and path length provide insights into the connectivity distance within the brain, while clustering coefficients reflect network segregation and its role in processing information within specialized brain modules [31]. A higher clustering coefficient indicates a greater tendency for the brain to perform specialized processes within these network modules [32]. Surprisingly, we found no significant differences in network segregation between patients with PNFA-FTD and HCs, and only the segregation of speech production network was found in PNFA-FTD according to previous studies [22]. Collectively, these findings suggest that the disruptions in global network topology are influenced, at least in part, by alterations in specific module-level integration and specialization. This novel insight sheds light on the role of specific modular changes in contributing to global network dysfunctions observed across different subtypes of FTD.

Consistent with prior research [20], our study identified substantial dysconnectivity within the SUB modules in both BV-FTD and SV-FTD, characterized by an increased number of intermodular connections involving SUB-SSN, SUB-VAN, and SUB-DMN, when compared to HC and individuals with PNFA-FTD. To delve into the details, BV-FTD displayed a notable decrease in intra-SUB connections but exhibited an increase in intermodular connections specifically between SUB and DMN compared to HC. Conversely, SV-FTD exhibited significantly higher interconnections between SUB and other brain modules, distinguishing it from both HC and PNFA-FTD. While FTD is traditionally associated with predominant atrophy in the frontal and temporal lobes, our findings underscore the critical role of subcortical structures in the progression of FTD, even in its presymptomatic stages [33, 34]. The SUB modules encompass various regions such as the amygdala, hippocampus, basal ganglia, and thalamus, all of which play pivotal roles in multiple cognitive and behavioral functions. The results from our correlation analysis align with these functional roles, demonstrating that the segregation of SUB is correlated with general cognition (CDR_total), behavioral control (CDR_behaviors), episodic verbal learning (CLVT), and functional activities (FAQ). More specifically, the amygdala, a limbic structure within the SUB, is implicated in emotional processing and reward learning [35]. Both BV-FTD and SV-FTD exhibit atrophy in the amygdala, a finding consistent with their shared behavioral features, including a lack of insight, impaired personal and social conduct, and disinhibition [7, 36]. The hippocampus, central to memory functions, is also vital for emotional regulation and the evaluation of facial expressions [37]. Emotional dysregulation is a core symptom of BV-FTD, while SV-FTD is characterized by deficits in facial emotion discrimination [35]. Previous meta-analyses have confirmed decreased gray matter volume in the amygdala and hippocampus in BV-FTD [38]. Additionally, graph theory measurements have shown that the severity of semantic deficits in SV-FTD is significantly correlated with the degree centrality values of the left anterior hippocampus [39]. The thalamus and basal ganglia, including structures like the putamen, caudate, globus pallidus, and nucleus accumbens, are highly complex and intricately connected to various brain regions. Dysconnectivity in the nucleus accumbens and caudate, which are regions associated with reward processing, has been reported in BV-FTD [35]. Overall, our findings highlight the substantial involvement of SUB modules in the progression of FTD, shedding light on the potential associations between SUB deterioration and FTD symptoms, an area that has not been extensively explored in prior research.

Apart from the SUB module, our study also revealed significant disorganization in the DMN, FPN, VAN, with increased numbers of connectivity of those modules in both BV-FTD and SV-FTD, compared to HCs. Disruptions in these networks, DMN, FPN, and VAN, have been described as a “triple model” observed in various diseases, including autism, schizophrenia, and AD [40]. This model suggests that the VAN acts as an interface between the DMN and FPN, regulating their competing inter-network activities and promoting appropriate behavioral responses [40, 41]. Our intra- and inter-module analyses identified altered connectivity patterns within the VAN, such as VAN-AN, VAN-DMN, and VAN-SUB. Moreover, we found correlations between the segregation of VAN and clinical variables, including general cognition (CDR_total), depression (GDS), episodic verbal learning (CLVT), and functional activities (FAQ). The VAN’s role in detecting and filtering relevant information from the environment, as well as its involvement in attention shifting and salient stimulus processing, has been consistently reported in prior studies [42, 43]. BV-FTD exhibited significant alterations in VAN connectivity compared to HCs and showed even more profound disruptions in VAN than AD, possibly due to the VAN’s association with psychiatric disorders [11, 20, 41]. Similarly, the VAN exhibited dysconnectivity in SV-FTD compared to HCs [44, 45]. The significant correlation between the MSI of VAN and cognitive decline aligns with previous findings suggesting that baseline measures of VAN connectivity, particularly in the left insula, may predict behavioral changes in patients with FTD, including BV-FTD and SV-FTD [43]. Additionally, a study by Chiong, et al. [42] revealed diminished influence from VAN to DMN during moral reasoning in BV-FTD, in contrast to the directed functional connectivity from VAN to DMN during moral reasoning observed in HC. The DMN, primarily active at rest and inactive during cognitive tasks, is associated with introspection, self-referential thinking, and mind-wandering [40, 42]. Our study identified alterations in DMN segregation in both BV-FTD and SV-FTD, highlighting the remarkable dysconnectivity observed in depression, characterized by excessive internal focus [40] Loss of empathy is a common feature in FTD, which shares similarities with depressive symptoms [46]. The FPN is responsible for cognitive control and goal-directed thinking [47]. We found a correlation between the MSI of FPN and behavioral control (CDR_behaviors), indicating that it is active during tasks requiring focused attention, decision-making, and problem-solving [40]. Notably, only BV-FTD exhibited altered FPN segregation, which may be linked to their uncontrolled behaviors. Previous research has shown that FPN connectivity in BV-FTD deteriorates with disease progression in longitudinal studies [48].

Generally, the significant degenerations observed in the VAN in BV-FTD and SV-FTD may contribute to the unbalanced segregation of the DMN in these subtypes and increased interconnections in the FPN associated with uncontrolled behaviors in BV-FTD [3].

These findings suggest the presence of a common underlying degeneration model shared between BV-FTD and SV-FTD. Prior studies have noted the similarities in neural network disruptions, particularly in the VAN and frontolimbic connectivity, between these two subtypes, reflecting the presence of similar behavioral abnormalities in BV-FTD and SV-FTD [36, 49].

### Transdiagnostic nodal to modulate the brain networks

Several key brain regions, including FG, OFC, left precuneus, and right insular emerged as crucial nodes influencing the segregation of brain networks in FTD subtypes. The OFC, vital for processing emotional signals, evaluating social and emotional context, and guiding appropriate behavioral responses, exhibited significant connectivity changes in BV-FTD [50]. This is in line with common alterations seen in psychiatric disorders, which share similarities with BV-FTD [51]. In our study, increased connectivity from the OFC to multiple brain networks, including DMN, FPN, AN, and VAN, was observed in BV-FTD. Structural atrophy [8] and reduced metabolism activity [52] have previously been reported in the OFC in BV-FTD. This increased connectivity to various brain networks may underlie the diverse behavioral abnormalities exhibited by BV-FTD individuals, such as emotional apathy, social reward learning deficits, impulsivity, poor judgment, and difficulty in understanding social cues [53]. Previous suggested the OFC degeneration might cause emotion recognition and interoceptive accuracy impairment in BV-FTD [54].

The insular cortex, a central brain region within the VAN, plays a pivotal role in functions such as salience detection, empathy, and language processing [55]. Our study identified significantly altered PC values in the insular cortex with decreased connectivity within VAN and between VAN and FPN in BV-FTD, SV-FTD, and PNFA-FTD. The insular cortex’s atrophy has been well-documented in all major FTD subtypes [8]. Notably, lateralization changes in the insular cortex appear to contribute to different FTD subtypes, with SV-FTD showing predominant atrophy in the left hemisphere and BV-FTD showing right hemisphere insular atrophy. These lateralized changes may result in reduced baseline autonomic output, often associated with the hallmark behavioral, language, and emotional disturbances observed in FTD [55, 56]. The shared involvement of the insular cortex in different FTD subtypes suggests a common underlying mechanism, possibly related to the spread of abnormal protein aggregates, such as tau or TDP-43, within neural networks [57, 58]. Understanding these common mechanisms could potentially lead to more effective diagnostic tools and targeted therapeutic interventions.

### Limitation and future work

This study has several limitations that warrant consideration and suggest directions for future research. Firstly, the relatively small sample size in this study may limit the generalizability of our findings. Expanding the sample size in future investigations would enhance the robustness and reliability of the results. Secondly, our use of functional networks derived from Yeo’s atlas may not fully capture the intricacies of speech abnormalities specific to PNFA-FTD. Future work should consider employing more tailored brain modules, like semantic appraisal network [59], speech production network [22], that better depict the neural correlates of PNFA-FTD, allowing for a more comprehensive understanding of its pathophysiology. Thirdly, our study focused solely on patients with the three major FTD subtypes. To gain a more holistic view of neurodegenerative diseases, it would be beneficial to include a broader spectrum of diagnostic groups, such as AD and Parkinson’s disease, in future studies. This approach would enable the exploration of both shared similarities and distinctive characteristics across these conditions, advancing our understanding of transdiagnostic and diagnosis-specific pathophysiological mechanisms. Finally, while our study offers insights into transdiagnostic and diagnosis-specific alterations in modular brain networks, the practical utility of these findings in developing effective psychiatric treatment strategies remains to be fully elucidated. Longitudinal studies are needed to assess the value of these alterations for predicting the onset of disorders at the prodromal stage and monitoring therapeutic progress. Such research would be invaluable in the development of more targeted and efficient treatments for neurodegenerative diseases and related psychiatric conditions.

## Conclusion

In summary, our high-resolution connectomic analysis revealed distinctive patterns of modular brain network alterations in BV-FTD, SV-FTD, and PNFA-FTD. Transdiagnostic findings suggest that BV-FTD and SV-FTD exhibit overall disruptions in network organization, which may be linked to their common behavioral abnormalities. Changes in the SUB, DMN, FPN, and VAN are observed, with implications for cognition and behavior. Additionally, specific nodes like the OFC, FG, precuneus, and insular cortex play pivotal roles in modulating network segregation. Diagnosis-specific findings indicate that BV-FTD exhibits broader modular changes compared to other subtypes. These insights into modular brain network alterations offer a foundation for more precise diagnostics and targeted therapeutic interventions, opening new avenues for understanding the complex nature of Frontotemporal Dementia and related disorders.

