## Supporting_information for "Transdiagnostic brain module dysfunctions across sub-types of frontotemporal dementia: a connectome-based investigation"

Table 1, the Significant between-group difference in modular measures, controlled by age and gender.

|  | **Group effect** | ***Post hoc*** **pairwise analyses** | | | | | |
| --- | --- | --- | --- | --- | --- | --- | --- |
|  |  | **BV-FTD vs. HC** | **SV-FTD vs. HC** | **PNFA-FTD vs. HC** | **BV-FTD vs. SV-FTD** | **BV-FTD vs. PNFA-FTD** | **SV-FTD vs. PNFA-FTD** |
| **Module segregation index** | |  |  |  |  |  |  |
| SUB | **F=9.5464, *p*<.0001(<.0001)** | **T=-5.170, *p<*.0001(<.0001)** | **T=-3.810, *p*=.0002(.001)** | T=.005, *p*=.996(1.0) | T=-.770, *p*=.044(.0868) | **T=-3.829, *p*=.0002(.001)** | **T=-2.982, *p*=.003(.017)** |
| DMN | **F=10.062, *p<*.0001(<.0001)** | **T=-3.545, *p*=.0007(.004)** | **T=-4.515, *p<*.0001(.001)** | T=.254, *p*=.8(.99) | T=1.216, *p*=.225(.617) | **T=-2.777, *p*=.006(.03)** | **T=-3.741, *p*=.0002(.001)** |
| FPN | **F=9.158, *p*<.0001(<.0001)** | **T=-5.372, *p<*.0001(<.0001)** | T=-1.43, *p*=.0154(.482) | T=-.555, *p*=.96(1.0) | **T=-3.022, *p*=.003(.015)** | **T=-3.926, *p*=.0001(.0007)** | T=-1.071, *p*=.285(.707) |
| VAN | **F=13.85, *p*<.0001(<.0001)** | **T=-5.372, *p<*.0001(<.0001)** | **T=-4.082, *p*=.0001(.0004)** | T=-1.811, *p*=.071(.271) | T=-0.692, *p*=.489(.09) | T=-2.395, *p*=.017(.08) | T=-.167, *p*=.09(.338) |
| **Intramodular connections** | |  |  |  |  |  |  |
| SSN-SSN | **F=4.978, *p*=.0024(.01)** | **T=-4.065, *p=*.0001(.0004)** | T=-1.441, *p*=.15(.475) | T=-0.459, *p*=.511(.912) | T=-1.972, *p*=.05(.202) | T=-2.433, *p*=.016(.074) | T=-.575, *p*=.566(.939) |
| VAN-VAN | **F=3.97, *p*=.0088(.0288)** | **T=-3.193, *p=*.0017(.009)** | T=-.965, *p*=.336(.78) | T=-.879, *p*=.381(.816) | T=-1.695, *p*=.092(.369) | T=-1.595, *p*=.112(.384) | T=-.02, *p*=.984(1) |
| **Intermodular connections** | |  |  |  |  |  |  |
| VN-SSN | **F=9.761, *p<*.0001(<.0001)** | **T=-4.815, *p<*.0001(<.0001)** | T=-1.501, *p*=.135(.439) | **T=-3.29, *p*=.001(.007)** | T=-2.51, *p*=.013(.06) | T=.692, *p*=.49(.9) | T=1.575, *p*=.117(.395) |
| SSN-AN | **F=4.128, *p*=.0073(.0262)** | **T=3.016, *p=*.0029(.015)** | **T=2.937, *p*=.004(.019)** | T=.062, *p*=.959(1.0) | T=.179, *p*=.857(.998) | T=2.285, *p*=.02(.10) | T=.067, *p*=.471(.565) |
| SSN-SUB | **F=5.928, *p*=.0007(.008)** | **T=3.092, *p*=.002(.010)** | T=1.122, *p*=.135(.203) | T=.276, *p*=.382(.203) | T=1.755, *p*=.039(.078) | T=2.013, *p*=.018(.053) | T=2.347, *p*=.20(.091) |
| DAN-FPN | **F=4.665, *p*=.004(.014)** | **T=-3.8, *p*=.0002(.0011)** | T=-1.424, *p*=.156(.486) | T=-.76, *p*=.448(.872) | T=-1.776, *p*=.077(.288) | T=-2.149, *p*=.03(.14) | T=-.478, *p*=.633(.964) |
| VAN-AN | **F=5.72, *p*=.0009(.0008)** | **T=3.827, *p*=.0002(.001)** | **T=3.04, *p*=.003(.014)** | T=-.198, *p*=.0843(.997) | T=.376, *p*=.707(.981) | **T=3.004, *p*=.003(.016)** | T=2.542, *p*=.011(.059) |
| VAN-DMN | **F=4.91, *p*=.003(.013)** | **T=3.098, *p*=.002(.011)** | **T=2.803, *p*=.006(.028)** | T=1.007, *p*=.315(.754) | T=.004, *p*=.997(1.0) | T=1.413, *p*=.159(.493) | T=1.349, *p*=.179(.53) |
| VAN-SUB | **F=5.059, *p*=.002(.013)** | **T=1.757, *p=*.081(.297)** | **T=3.908, *p*=.0001(.0007)** | T=.364, *p*=.715(.983) | T=-2.035, *p*=.043(.179) | T=.982, *p*=.327(.760) | **T=2.75, *p*=.006(.038)** |
| AN-FPN | **F=6.318, *p*=.0004(.007)** | **T=3.575, *p=*.0004(.003)** | T=2.073, *p*=.04(.166) | T=-.663, *p*=.508(.901) | T=1.027, *p*=.306(.734) | **T=3.224, *p*=.0015(.008)** | T=2.174, *p*=.03(.134) |
| DMN-SUB | **F=4.666, *p*=.0036(.014)** | T=2.555, *p*=.011(.055) | **T=2.824, *p*=.005(.027)** | T=1.106, *p*=.27(.687) | T=-.447, *p*=.656(.97) | **T=-2.855, *p*=.005(.024)** | T=3.131, *p*=.002(.108) |
| **Nodal participation coefficient** | |  |  |  |  |  |  |
| L. FG | **F1=16.296, F2=9.800(62)** | **T=-6.516, *p*<.0001(<.0001)** | T=.113, *p*=.91(1.0) | T=-.475, *p*=.635(.964) | **T=-4.406, *p<*.0001(.0001)** | **T=-5.289, *p<*.0001(.0001)** | T=.485, *p*=.628(.962) |
| L. OFC | **F1=9.487, F2=7.582(55)** | **T=-5.023, *p*<.0001(<.0001)** | T=-1.674, *p*=.10(.34) | T=-1.051, *p*=.294(.719) | **T=-3.077, *p*=.002(.012)** | T=-2.255, *p*=.025(.113) | T=.577, *p*=.564(.938) |
| R. FG | **F1=14.383, F2=7.749(52)** | **T=-5.686, *p*<.0001(<.0001)** | T=.898, *p*=.37(.805) | **T=-3.432, *p*=.0007(.004)** | **T=-3.73, *p*=.0002(.001)** | T=-1.212, *p*=.227(.602) | T=2.16, *p*=.03(.137) |
| L. precuneus | **F1=10.166, F2=8.576(43)** | **T=-4.821, *p*<.0001(<.0001)** | T=1.01, *p*=.036(.155) | T=-1.674, *p=*.096(34) | **T=-4.735, *p*<.0001(<.0001)** | T=-2.106, *p*=.037(.155) | T=-2.195, *p*=.029(.128) |
| R. insular | **F1=9.117, F2=8.097(39)** | **T=3.763, *p=*.0002(.0013)** | **T=3.364, *p=*.0009(.0051)** | **T=3.502, *p=*.0006(.0032)** | T=.041, *p=*.967(1.0) | T=-.272, *p=*.786(.993) | T=.297, *p=*.767(.991) |

Data are presented as F/T-statistics and uncorrected *p*-values (FDR-corrected *p*-values are shown in parentheses). Values presented in bold indicate an FDR-corrected significance level of *p*<0.05. Notably, F1 for the nodal participation coefficient are presented group difference and *F2* for the nodal participation coefficient are presented as the *F* value for the peak within the cluster (cluster size). HC, healthy control; BV-FTD, behavioral variant frontotemporal dementia; SV-FTD, semantic variant frontotemporal dementia; PNFA-FTD, progressive non-fluent aphasia frontotemporal dementia; SUB: subcortical module; DMN: default mode network; FPN: frontoparietal network; VAN: ventral attention network; AN: affective network; DAN: dorsal attention network; SSN: sensorimotor network; VN: visual network; FG: fusion gyrus; OFC: orbitofrontal cortex; L, left; R, right.
